# Lineage-specific control of convergent differentiation by a Forkhead repressor

**DOI:** 10.1101/758508

**Authors:** Karolina Mizeracka, Julia M. Rogers, Jonathan D. Rumley, Shai Shaham, Martha L. Bulyk, John I. Murray, Maxwell G. Heiman

## Abstract

During convergent differentiation, multiple developmental lineages produce a highly similar or identical cell type. However, few molecular players that drive convergent differentiation are known. Here, we show that the *C. elegans* Forkhead transcription factor UNC-130 is required in only one of three convergent lineages that produce the same glial cell type. UNC-130 acts transiently as a repressor in progenitors and newly-born terminal cells to allow the proper specification of cells related by lineage rather than by cell type or function. Specification defects correlate with UNC-130:DNA binding, and UNC-130 can be functionally replaced by its human homolog, the neural crest lineage determinant FoxD3. We propose that, in contrast to terminal selectors that activate cell-type specific transcriptional programs in terminally differentiating cells, UNC-130 acts early and specifically in one convergent lineage to produce a cell type that also arises from molecularly distinct progenitors in other lineages.

## INTRODUCTION

Development is the story of how a single cell divides to give rise to lineages that produce every cell type in the body. The standard framework for understanding this process is that cell lineages branch to produce increasingly divergent cell states, with each cell type produced exclusively by a single branch (Fig. 1A). An exception to this paradigm is convergent differentiation, a phenomenon in which multiple lineages produce identical or highly similar cell types (Fig. 1B). For example, mesoderm and neural crest lineages both produce the same type of heart cell (Dupin et al., 2018; Keyte & Hutson, 2012), and embryonic and extra-embryonic lineages both produce gut endoderm (Kwon et al., 2008). With the advent of single-cell RNA profiling coupled to lineage tracing, it is now appreciated that convergent differentiation is surprisingly prevalent in vertebrates (Chan et al., 2019; Liu et al., 2019; McKenna & Gagnon, 2019; Wagner et al., 2018). However, the molecular players that drive convergent differentiation remain unclear.

**Fig. 1.**
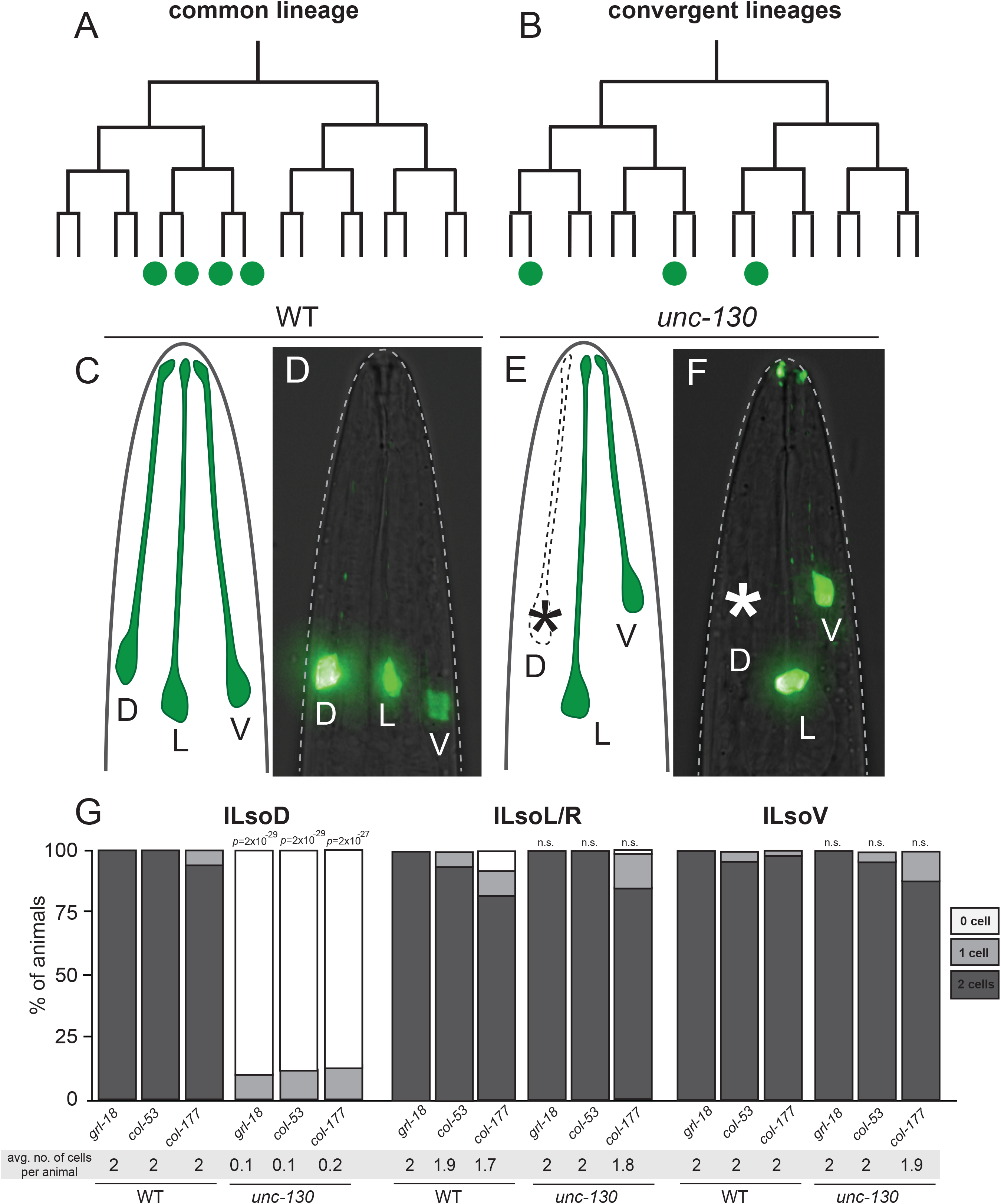
UNC-130 is required in one of three convergent lineages. Schematic of a common lineage (A), in which shared progenitors give rise to a unique cell type and a convergent lineage (B), in which distinct progenitors in divergent lineages produce a unique cell type. Lateral views and schematics of wild-type (C, D) and *unc-130* mutant (E, F) animals expressing ILso marker, *grl-18*pro:YFP. Asterisk denotes missing cells. D – dorsal, L – lateral, V – ventral. (G) Percentage of wild-type and *unc-130* mutant animals expressing ILso markers *grl-18*, *col-53*, and *col-177* in zero, one, or two dorsal (ILsoD), lateral (ILsoL/R) and ventral (ILsoV) glia. Average number of cells expressing each marker per animal is listed under each condition. n = 50 animals per marker and genotype. *p*-values were calculated by Fisher’s Exact test.

Importantly, in *C. elegans*, convergent differentiation has been appreciated for decades, ever since the complete developmental lineage was mapped (Sulston et al., 1983). A number of cell types that are present in symmetric anatomical regions – such as the dorsal and ventral equivalents of a given cell type – are produced through convergent differentiation by multipotent sublineages, which we will refer to as convergent lineages. A key question to understanding convergent differentiation is to determine if progenitors always follow the same path to produce a given cell type – that is, are shared or lineage-specific transcriptional trajectories employed across lineages that produce the same cell type? Intriguingly, several *C. elegans* mutants affect only a subset of convergently-derived cells: for example *mls-2* mutations affect ventral, but not dorsal CEP sheath glia; *lin-32* mutations affect dorsal, but not ventral CEP neurons; and *ceh-10* mutations affect dorsal, but not lateral or ventral RME neurons (Doitsidou et al., 2010; Forrester et al., 1998; Rojo Romanos et al., 2017; Yoshimura et al., 2008). These findings suggest that convergent lineages use distinct transcriptional trajectories to specify the same cell type. However, the mechanism by which transcription factors act across lineages to mediate convergent differentiation is not understood.

Most *C. elegans* glia are located in symmetric groups of sense organs in the head, called the inner labial (IL), outer labial (OL), cephalic (CEP), and amphid (AM) sensilla. Each organ contains exactly two glial cell types - the sheath and socket. We focused on the development of the IL socket glia which are six-fold radially symmetric, such that there is a dorsal, lateral, and ventral pair of cells (ILsoD, ILsoL/R, ILsoV, respectively) (Mizeracka & Heiman, 2015). We recently identified specific markers for ILso glia and determined a role for them during dendrite extension of associated sensory neurons (Cebul et al., 2020). ILso glia develop via convergent differentiation - all three pairs of cells arise from distinct lineages that diverge at early stages of embryogenesis (Sulston et al., 1983). Importantly, all three pairs of ILso glia express the same reporter genes and appear as a uniform cluster in single-cell profiling experiments (Packer et al., 2019). In contrast, the three lineage-specific pairs of ILso parent cells cluster separately, suggesting that these progenitors are molecularly distinct. In single cell RNA-sequencing studies, the ILso parent cells could not be identifiably linked to their terminal progeny, and thus were interpreted to develop through a “discontinuous” transcriptional trajectory (Packer et al., 2019). This may be because the transcriptomes of ILso parent cells change quickly after their terminal divisions as they undergo convergent differentiation, thereby making it difficult to identify factors that are important for this process.

Here, we find a lineage-specific role for the conserved Forkhead transcription factor UNC-130 during the specification of a convergently-derived glial cell type in *C. elegans*. We show that, during embryogenesis, UNC-130 is required for the specification of ILso cells that are derived from one convergent lineage, but is dispensable for the production of ILso cells derived from other lineages. Furthermore, consistent with previous work (Sarafi-Reinach & Sengupta, 2000), mutations in *unc-130* perturb the specification of several cell types that are related by lineage, but not by function. We find that UNC-130 is a transiently expressed transcriptional repressor that acts at the time of birth of the ILsoD glia. The vertebrate homolog of UNC-130, FoxD3, also acts in a lineage-specific manner and is required for the specification of neural crest-derived cell types (Kos et al., 2001; Lister et al., 2006; Stewart et al., 2006). Intriguingly, we find that UNC-130 and FoxD3 share molecular features, including their preferred binding sites, and FoxD3 can functionally replace UNC-130. Lineage-specific regulatory factors like UNC-130/FoxD3 may represent an evolutionarily ancient mechanism that enables molecularly distinct progenitors in different lineages to produce the same cell type.

## RESULTS

### *unc-130* mutants display defects in convergent differentiation

To identify genes controlling ILso glial fate, we performed a chemical mutagenesis screen and isolated a mutant strain in which one pair of ILso cells was missing marker expression (see Methods). Genetic mapping and sequencing revealed a causal mutation in the *unc-130* gene, which encodes a conserved Forkhead transcription factor. We obtained the reference allele for this gene, the deletion strain *ev505*, and characterized sense organ perturbations in detail (Nash et al., 2000). In previous work, we identified the gene *grl-18*, which encodes for a poorly characterized protein that contains a ground-like domain, as well as *col-53* and *col-177*, which both encode collagens, as highly specific markers for ILso glia (Cebul et al., 2020; Fung et al., 2020). Using these markers, we find that 100% of *unc-130* mutants lost expression of all three known ILso markers (*grl-18* pro, *col-53* pro, and *col-177* pro) specifically in one or both dorsal ILso glia (ILsoD), but not in lateral (ILsoL/R) or ventral (ILsoV) glia (Fig. 1C-G, Table S1; see Methods). ILso glia are associated in sense organs with two neurons called IL1 and IL2, and the IL sheath glial cell (ILsh). We find that loss of marker expression was specific to ILsoD glia – *unc-130* mutants occasionally have extra IL1 neurons and wild-type numbers of IL2 neurons (Table S2). Lack of specific reporters precluded examination of IL sheath glia.

Loss of marker expression in ILsoD glia in *unc-130* mutants could be due to defects in the cell division pattern of the lineage that normally produces these cells. For example, the progenitor cell could fail to divide, or the presumptive ILsoD cells could undergo cell death. To test these possibilities, we performed lineaging analysis on *unc-130* mutants to track all cell divisions in this sublineage. In 8/8 *unc-130* mutant embryos, we found that ILsoD progenitors divide to produce presumptive ILsoD glia and their sister cell, the skin cell hyp3 (Fig. S1). However, we found that the timing of this cell division was delayed compared to wild-type embryos. Cell cycle lengths of the other ILso progenitor cells were also slightly delayed in *unc-130* mutants, although these cells were not as strongly affected as ILsoD progenitors (Fig. S1). These results provide evidence that the putative ILsoD cells are born at approximately the right time in *unc-130* mutants, although the ILsoD progenitor may be abnormal as reflected by its delayed cell division. We conclude that loss of marker expression in ILsoD glia in *unc-130* mutants is likely due to a loss of identity, rather than changes in the division patterns of the ILsoD lineage.

### UNC-130 is required to produce cell types that are related by lineage, not by identity or function

We wanted to determine whether UNC-130 acts in ILsoD glia specifically or more broadly at the level of a sublineage. Previous work showed that UNC-130 is required for the specification of three sensory neurons (AWA, ASG, and ASI) that are produced by an exclusively neuronal sublineage that is most closely related to the ILsoD sublineage (Fig. 2A) (Sarafi-Reinach and Sengupta, 2000). This left open the possibility that UNC-130 is specific to neuronal fates. In contrast, the ILsoD sublineage is multipotent, giving rise to two types of glia, a neuron, and a skin cell (Fig. 2A) (Sulston et al., 1983).

**Fig. 2.**
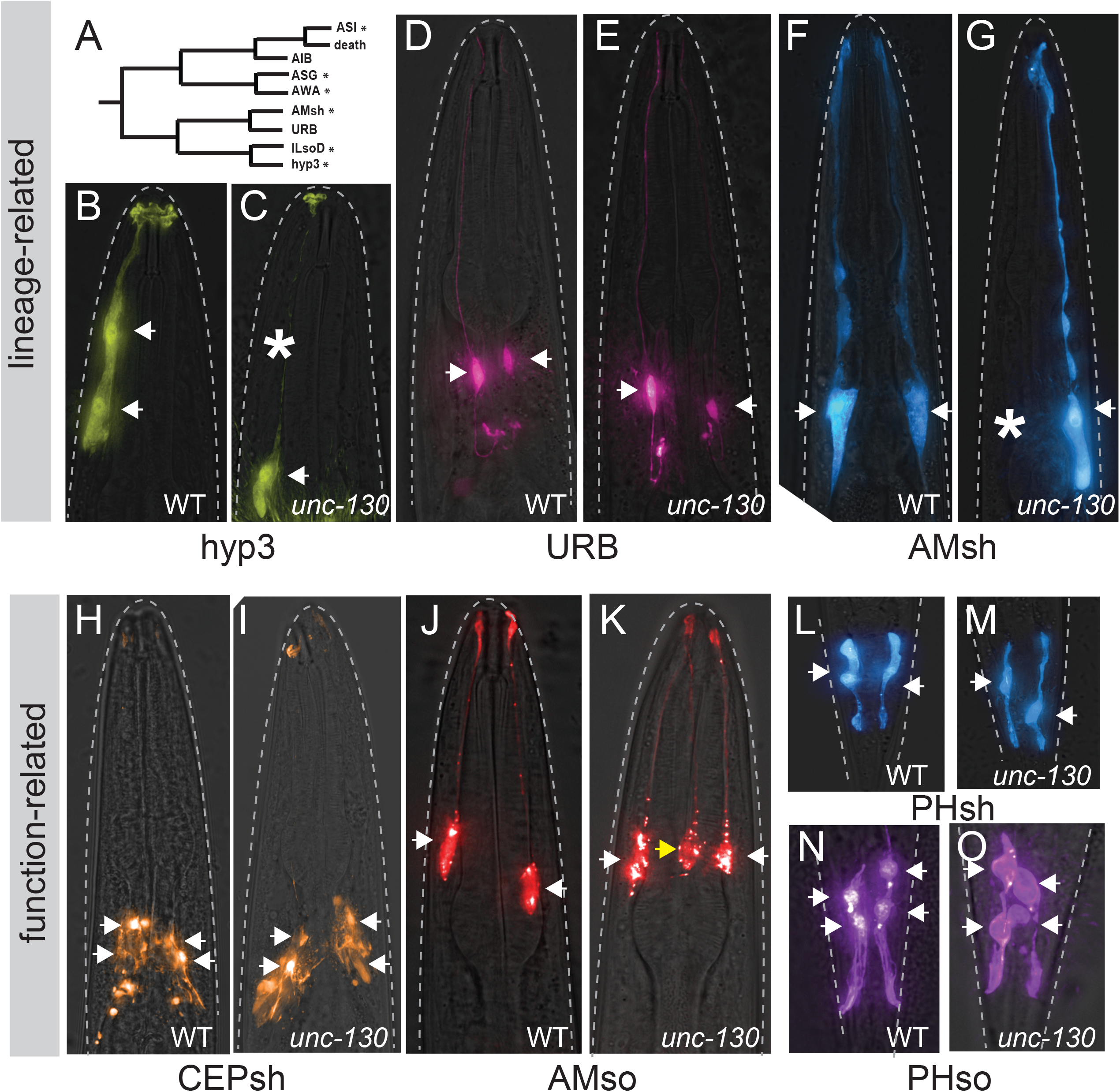
UNC-130 is required for the specification of lineage-related cell types. (A) Schematic of ILsoD sublineage. Asterisks denote cell types whose specification is affected in *unc-130* mutants. Wild-type (B) and *unc-130* mutant (C) animals expressing hyp3 marker, *ceh-10*pro:GFP. Asterisk denotes missing cell. Wild-type (D) and *unc-130* mutant (E) animals expressing URB marker, *nlp-6*pro:GFP. Wild-type (F) and *unc-130* mutant (G) animals expressing AMsh marker, *F16F9.3*pro:mCherry. Asterisk denotes missing cell. Wild-type (H) and *unc-130* mutant (I) animals expressing CEPsh marker, *hlh-17*pro:GFP. Wild-type (J) and *unc-130* mutant (K) animals expressing AMso marker, *grl-2*pro:mCherry. Wild-type (L) and *unc-130* mutant (M) animals expressing PHsh marker, *F16F9.3*pro:mCherry. Wild-type (N) and *unc-130* mutant (O) animals expressing PHso marker, *grl-2*pro:YFP. Arrows denote cell bodies in all images. Yellow arrow denotes extra AMso cell.

To systematically determine the requirement for UNC-130 in the ILsoD sublineage, we examined markers of each cell type: the hyp3 skin cell, the URB neuron, and the amphid sheath (AMsh) glial cell (Fig. 2A). In wild-type animals, there are two hyp3 cells whose cell bodies are located dorsally and fuse to form a syncytium. By mining single-cell transcriptome data, we identified *ceh-10*pro:GFP as a hyp3-specific marker (Packer et al., 2019, Fig. 2B). *ceh-10* encodes a conserved ortholog of the transcription factor CHX10 that is also expressed in unrelated neurons (Altun-Gultekin et al., 2001; Forrester et al., 1998). We found that in 30% of *unc-130* mutants, one or sometimes both hyp3 cells lacked marker expression (Fig. 2B,C, Table S3). Next, we used the same approach to identify a unique marker for the sensory neuron URB, the gene *nlp-6*, which encodes an uncharacterized neuropeptide (Packer et al, 2019, Fig. 2D). URB neurons form a bilateral pair on either side of the head (Fig. 2D). Specification of URB appeared unaffected in *unc-130* mutant animals, as these cells exhibited unchanged expression of *nlp-6*pro:GFP (Fig. 2E, Table S3). In contrast, specification of AMsh glial cells, which are sister cells of URB and are found in the bilaterally symmetric amphid sense organ, was affected, with 38% of *unc-130* mutants failing to express an AMsh-specific marker, *F16F9.3* which encodes a secreted peptide, in one or occasionally both cells (Fig. 2F,G, Table S3). Loss of *F16F9.3* expression was highly correlated with loss of two other AMsh glia-specific markers, *T02B11.3*pro:GFP and *F53F4.13*pro:GFP (Fig. S2A). Through transcriptional profiling experiments, these three markers have been determined to be highly specific for the AMsh, but their functions remain unknown (Bacaj et al., 2008; Fung et al., 2020; Wallace et al., 2016). Thus, we find that the specification of non-neuronal cell types that are derived from the ILsoD sublineage is affected in the absence of UNC-130.

Cell types that share an identity, for example neurons that express the same neurotransmitter, often express the same combination of transcription factors. To test whether UNC-130 affects cell types that are functionally related to ILso, we examined markers for other glial cell types. We found no defects in the specification of the four cephalic sheath glia (CEPsh); two phasmid sheath glia (PHsh), which are the functional equivalent of the AMsh glia in the tail and express many of the same markers; or four phasmid socket glia (PHso) (Fig. 2H,I, L-O, Table S3) (Fung et al., 2020; McMiller & Johnson, 2005; Yoshimura et al., 2008).

Surprisingly, we observed extra amphid socket (AMso) glia in *unc-130* mutants, with 42% of mutant animals expressing the AMso-specific marker, *grl-2,* in an extra cell (Fig. 2J, K, Table S3) (Hao et al., 2006). *grl-2* encodes an uncharacterized protein that includes a ground-like domain (Hao et al., 2006). We found that expression of *grl-2* in the extra cell was highly correlated with expression of two other AMso-specific markers, *lin-48,* which is the *C. elegans* ortholog of Ovo-like transcription factors in vertebrates, and *itr-1,* which encodes for an inositol triphosphate receptor (Fig. S2C) (Fung et al., 2020; Heiman & Shaham, 2009; Johnson et al., 2001; Low et al., 2019). Because *lin-48* encodes for a highly conserved transcription factor that is expressed specifically in AMso, it is possible that it also plays a role in the specification of these cells. However, we found that mutations in *lin-48* do not affect endogenous or ectopic AMso specification (Fig. S2D). We considered the possibility that the extra AMso glia could reflect duplication of the entire sublineage that normally gives rise to the AMso. Therefore, we examined specification of the CEM neuron, which is the sister cell of AMso glia in males, in wild-type and *unc-130* animals. No extra CEM neurons were observed in mutant males, providing evidence that ectopic AMso do not arise from a sublineage duplication (Fig. S2B). Finally, we noted that, in contrast to endogenous AMso cells, which are positioned laterally, extra AMso cells were located dorsally, consistent with the position of the missing ILsoD and hyp3 cells (Fig. S2E). This raises the possibility that the extra AMso cells might arise by mis-specification of ILsoD, hyp3, or both, although due to the timing of marker expression in the embryo we were unable to test this directly.

Taken together, these observations suggest that in the absence of UNC-130, several cell types – including neurons, glia, and skin – fail to be specified or possibly take on alternative fates. Importantly, defects in fate specification in *unc-130* mutants are not solely related to a particular cell’s identity or function, but rather to cell types that share a lineage origin.

### UNC-130 acts in lineage-specific progenitors and newly-born precursor cells

Due to its broad effects on several cell types arising from the same sublineage, we reasoned that, unlike terminal selectors, UNC-130 might act earlier in development. Indeed, previous work showed that UNC-130 is expressed in the immediate progenitors but not in the terminal cells of affected sensory neurons (Sarafi-Reinach & Sengupta, 2000). We examined a translational reporter strain in which UNC-130 was fused with GFP and acquired a time course of images during *C. elegans* embryogenesis (Sarov et al., 2006). Consistent with previous work and our lineaging experiments, UNC-130:GFP is brightly expressed at early stages of development (∼300 min to 500 min), and starts to wane as animals near hatching at 3-fold stages (550+ min) (Fig. 3A-G). In contrast, expression of the IL socket-specific marker, *grl-18*, is not detectable at early stages, starts to be expressed in 2-fold embryos (500 min), and stays on into adulthood (Fig. 3H-N). These results provide evidence that *unc-130* is expressed early in development, in progenitor cells and presumptive ILso cells after they are born, but its expression is downregulated as these cells differentiate.

**Fig. 3.**
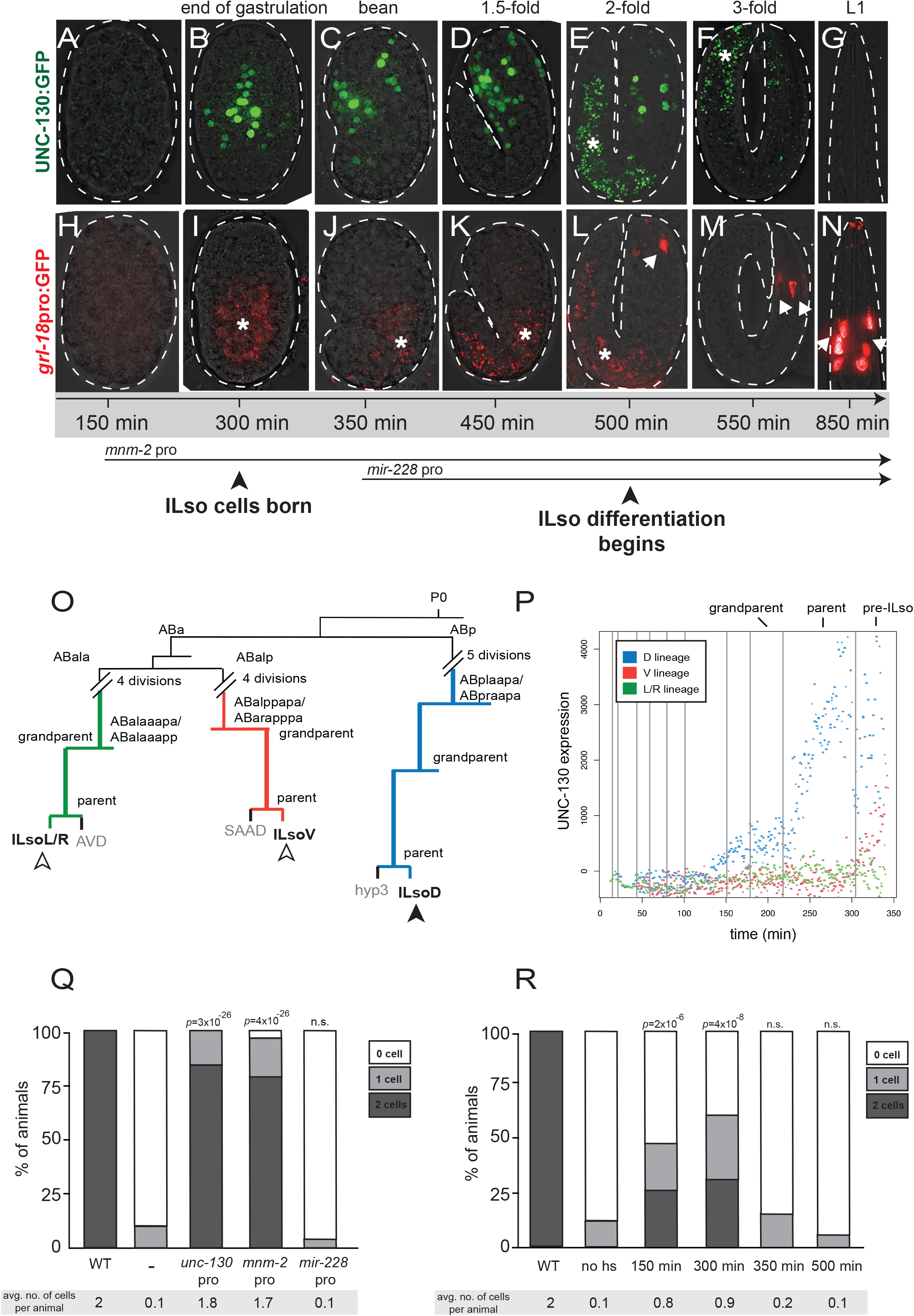
UNC-130 is expressed in a restricted lineage before the onset of differentiation. (A-G) Time course of embryos expressing UNC-130:GFP. (H-N) Time course of embryos expressing *grl-18*pro:GFP (pseudo-colored red). Asterisk denotes gut autofluorescence. Arrows denote *grl-18*+ ILso cells. (O) Lineage diagram of a subset of embryonic cell divisions derived from the AB blastomere that give rise to ILsoD (blue), ILsoV (red), ILsoL/R (green) glia. (P) Levels of UNC-130:GFP signal in ILsoD (blue), ILsoV (red), and ILsoL/R (green) lineages over developmental time. Vertical lines denote cell divisions. Subtraction of background fluorescence signal results in negative values for some cells. (Q) Heterologous promoters were used to drive expression of *unc-130* in the *unc-130* mutant strain and extent of rescue was assessed. Timing of *mnm-2*pro and *mir-228*pro expression is marked in (A). Percentage of animals expressing *grl-18*pro:YFP in zero, one, or two ILsoD glia in each condition. Average number of cells marked per animal is listed under each condition. n = 50 animals per condition. *p*-values were calculated by Fisher’s Exact test. (R) *unc-130* mutant embryos expressing hsp:*unc-130* were heatshocked at different time points and extent of rescue was assessed. Percentage of animals expressing *grl-18*pro:YFP in zero, one, or two ILsoD glia in each condition. Average number of cells marked per animal is listed under each condition. n = 50 animals per condition. *p*-values were calculated by Fisher’s Exact test.

We wanted to understand why mutation of *unc-130* affects the specification of the ILsoD pair of glia, but not the equivalent lateral or ventral pairs. We noted that the lineages that give rise to the three pairs of ILso glia diverge at the 4- and 16-cell stages (Fig. 3O) (Sulston et al., 1983). Previous studies showed that *unc-130* expression commences midway through embryogenesis in an ABp-derived lineage that gives rise to dorsal – but not lateral or ventral – ILso glia (Murray et al., 2012; Sarafi-Reinach and Sengupta, 2000). To better define this expression pattern, we performed UNC-130 lineaging experiments and extended the period of data acquisition to capture the birth of the presumptive ILso cells. We found that *unc-130* is highly expressed in the ILsoD lineage and in the presumptive ILsoD cell after it is born, but not in the lineages that give rise to the ILsoL/R or ILsoV cells (Fig. 3P, S3). We did note low but increasing levels of expression in ILsoV glia after they are born. The significance of this is unclear, as ILsoV glia are specified normally in *unc-130* mutants. Taken together, these findings show that UNC-130 is expressed highly in the progenitor that gives rise to the ILsoD glia around the time when the cells are born.

To functionally address when UNC-130 is required for fate specification, we used early and late promoters to drive *unc-130* expression in *unc-130* mutants and determined the extent of rescue of glial specification defects by scoring for the presence of *grl-18*+ cells in late larvae or young adults. For early rescue, we used a regulatory sequence we identified upstream of *mnm-2*, a gene which, similarly to *unc-130,* is expressed early in the ILsoD sublineage but not in other lineages that normally express *unc-130* (Fig. S4) (Murray et al., 2012). For late rescue, we used a sequence upstream of *mir-228,* which is expressed in all glial cells shortly after they are born (Fig. S4) (Fung et al., 2020; Pierce et al., 2008). We find that early *unc-130* expression under the *mnm-2* promoter in *unc-130* mutants rescues specification of ILsoD glia almost as completely as expression driven by the *unc-130* promoter (Fig. 3Q). In contrast, late expression of *unc-130* driven by the *mir-228* promoter does not rescue specification defects (Fig. 3Q). Intriguingly, expression of *mir-228*pro:*unc-130* in *unc-130* mutants resulted in a mild enhancement of specification defects such that a larger percentage of animals lacked *grl-18* expression in ILsoD cells. This suggests that downregulation of *unc-130* in newly-born cells is important for fate specification.

In a complementary approach, we drove *unc-130* expression using a heat-shock inducible promoter in *unc-130* mutants to define a temporal window during which UNC-130 is required for fate specification. We found that heat-shock induction of UNC-130 at early stages of development resulted in a high lethality rate, therefore, we scored arrested embryos or larvae 12-18 hours after heat shock. Heat-shock induced expression of *unc-130* at ∼150 min (see Methods), before the birth of the ILsoD, resulted in moderate rescue with 48% of animals expressing *grl-18* in one or two ILsoD cells as compared to 12% of animals expressing *grl-18* in one ILsoD cell in no heat-shock controls (Fig. 3R). Heat-shock-induced expression of *unc-130* at approximately the birth of ILsoD glia (300 min) resulted in the strongest rescue of fate specification defects with 62% of animals expressing *grl-18* in one or two ILsoD glia (Fig. 3R). Heat-shock induction after ∼350 min, which is after the ILsoD cells are born but before they start to differentiate, or any other later time points, did not result in significant rescue of specification defects as compared to no heat-shock controls (Fig. 3R).

Together these results provide evidence that *unc-130* acts transiently as ILsoD progenitors are dividing to produce ILsoD glia and shortly after these cells are born, but it cannot rescue specification defects after this temporal window has passed. This temporal requirement for UNC-130 before the onset of differentiation suggests that rather than directly instructing cell fate, UNC-130 establishes competency for fate specification.

### UNC-130 functions as a repressor

The UNC-130 homolog FoxD3 acts as a repressor in several developmental contexts, in some cases recruiting the Groucho repressor complex through a conserved engrailed homology (eh1) domain (Ono et al., 2014; Yaklichkin, Steiner, et al., 2007). Paradoxically, FoxD3 also functions as a pioneer factor in embryonic stem cells and during early neural crest specification (Krishnakumar et al., 2016; Lukoseviciute et al., 2018). To determine whether UNC-130 acts as a repressor or an activator during convergent differentiation, we expressed synthetic constructs encoding the UNC-130 DNA-binding domain (DBD) fused to a canonical activator (VP64 – four copies of VP16 activator domain) or repressor (*Drosophila* Engrailed domain) under the *unc-130* promoter in an *unc-130* mutant strain and assessed rescue of glia specification defects by scoring for the presence of *grl-18*+ cells in late larvae or young adults (Fig. 4A,B). Expression of full-length UNC-130 fully rescued ILsoD glia defects (Fig. 4B). Expression of the UNC-130 DBD alone or a DBD-VP64 synthetic activator showed no rescue (Fig. 4B). In contrast, the synthetic repressor DBD-Engrailed substantially rescued ILsoD specification defects, suggesting that UNC-130 normally functions as a transcriptional repressor (Fig. 4B).

**Fig. 4.**
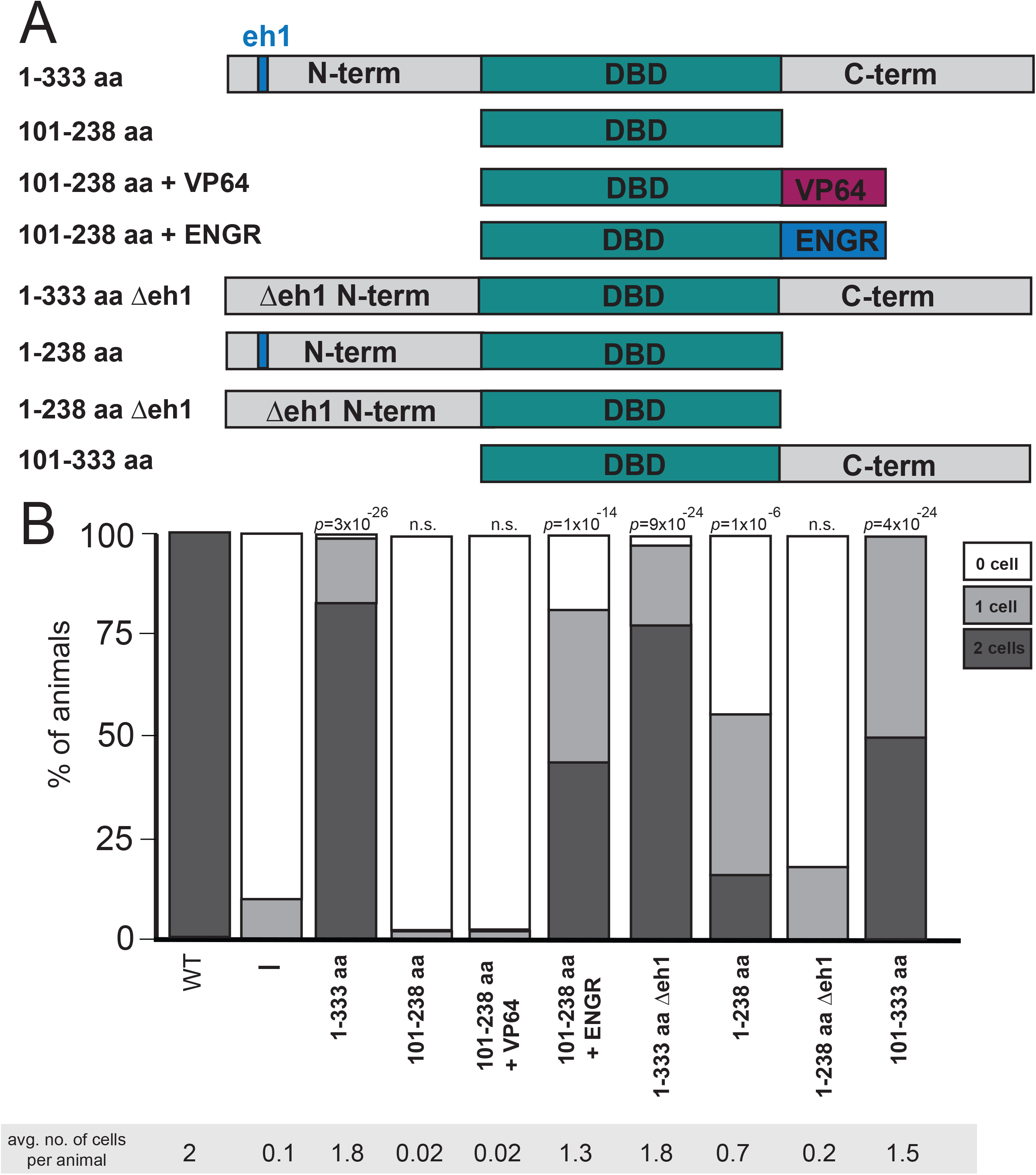
UNC-130 acts as a repressor to specify glial fates. (A) Schematic diagram of UNC-130 constructs used in rescue experiments to determine domain function. Full-length UNC-130 is 333 amino acids in length. (B) *unc-130* promoter was used to drive expression of individual constructs (schematized in A) in the *unc-130* mutant strain and extent of rescue was assessed. Percentage of animals expressing *grl-18*pro:YFP in zero, one, or two ILsoD glia in each condition. Average number of cells marked per animal is listed under each condition. n = 50 animals per condition. *p*-values were calculated by Fisher’s Exact test.

The vertebrate UNC-130 homolog, FoxD3, contains an eh1 motif in its carboxy-terminal domain that recruits the Groucho repressive complex. This region of UNC-130 is not conserved, but we identified a candidate eh1 sequence in the UNC-130 amino-terminal domain and tested whether it might act similarly (Yaklichkin, Vekker, et al., 2007) (Fig. 4A, S5, S6A). Surprisingly, deletion of the eh1 domain from full-length UNC-130 did not prevent rescue of specification defects, due to a redundant function of the carboxy-terminus as discussed below (Fig. 4, 1-333 aa Δeh1). We found that expression of a protein containing the UNC-130 amino-terminus and DBD partially rescues ILsoD defects, and the candidate eh1interaction motif is necessary for this rescuing function (Fig. 4, 1-238 aa and 1-238 aa Δeh1).

Interestingly, expression of the DBD with the carboxy-terminus also rescues all glial specification defects, which suggests it might contain previously unidentified repressor motifs as well (Fig. 4, 101-333 aa). Using luciferase assays in cultured mammalian cells, we found that N-term:GAL4DBD:C-term and GAL4DBD:C-term have reduced reporter activity compared to GAL4 DBD alone (Fig. S5B,C), consistent with the carboxy-terminus harboring repressive activity.

We independently tested the role of the Groucho complex by examining mutants in the sole ortholog of Groucho, *unc-37*, in *C. elegans*. *unc-37* mutants did not display any defects in ILso glial specification, which provides further evidence that UNC-130 harbors a redundant, Groucho-independent repressive motif (Fig. S5A). By contrast, we find that rescue with the N-terminus alone shows a strong dependence on *unc-37*, providing evidence that the N-terminal domain acts through recruitment of the Groucho repressive complex akin to vertebrate Forkhead factors (Fig. S5A).

Thus, we find that UNC-130 promotes ILsoD fate by acting as a transient transcriptional repressor at the time when these glial cells are produced. One possibility is that UNC-130 directly represses genes that would activate alternative fates, allowing presumptive ILsoD glia to follow the correct transcriptional trajectory.

### UNC-130 and the neural crest determinant FoxD3 share a conserved function

The UNC-130 DBD is highly conserved with its vertebrate homolog, FoxD3 (Fig. 5A, S6A), thus we wanted to determine if they share DNA-binding specificities. Different Forkhead transcription factors can bind to a consensus primary motif RYAAAYA (FkhP), a related secondary motif AHAACA (FkhS), or an unrelated alternate motif GACGC (FHL) (Nakagawa et al., 2013). To determine its binding specificity, we incubated recombinant UNC-130-DBD protein with universal protein binding microarrays (PBMs) containing all possible 10-mer double-stranded DNA sequences (Berger et al., 2006). We find that UNC-130-DBD preferentially binds [A/G][T/C]AAACA and AA[T/C]AACA sequences, variants of the primary and secondary Forkhead binding motifs, respectively, consistent with it behaving as a conserved Forkhead family member (Fig. 5B).

**Fig. 5.**
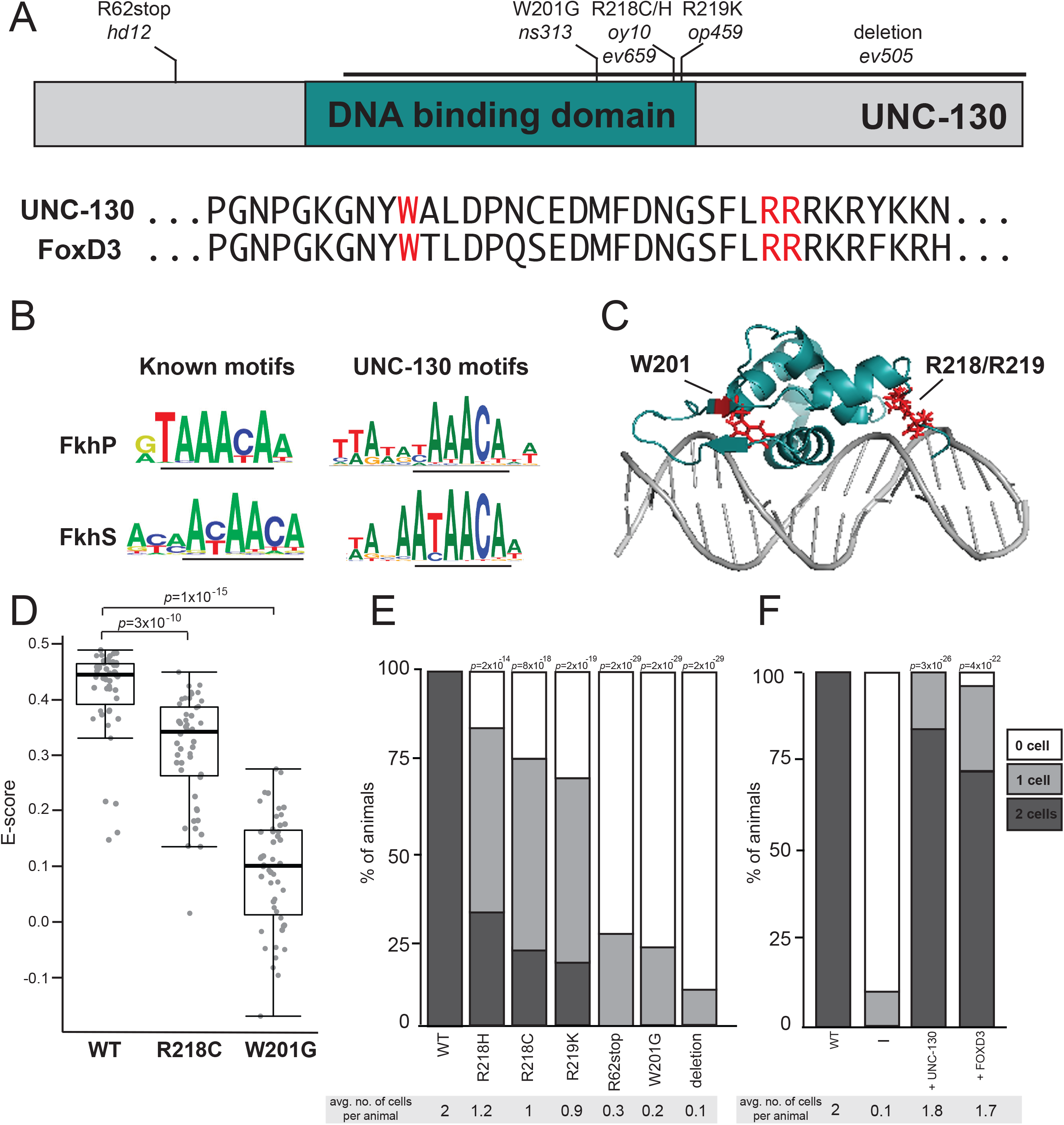
Severity of glial phenotypes correlate with UNC-130:DNA binding defects. (A) Schematic diagram of UNC-130 protein with DNA-binding domain (turquoise) highlighted. Location of point mutations and deletion are indicated. Alignment of UNC-130 and FoxD3 DBD with point mutations highlighted in red. (B) Logos of vertebrate primary (FkhP, top) and secondary (FkhS, bottom) Forkhead motifs and UNC-130 preferred DNA sequences that resemble primary (top) and secondary (bottom) motifs as determined by this study. Core motifs are underlined. (C) Structure of FoxD3 DBD (turquoise) interacting with DNA (gray) (Protein Data Bank ID code 2HDC, (Jin et al., 1999)). W201, R218, and R219 residues are highlighted in red. (D) Scatter plot of E-scores for 8-mer DNA sequences matching [A/G][C/T]AAACA or AA[C/T]AACA from protein binding microarray assays of wild-type, R218C, and W201G mutant UNC-130 proteins. Black lines represent population median; top and bottom of boxes are 25^th^ and 75^th^ percentiles, respectively; and top and bottom of whiskers are either most extreme point or 1.5x the interquartile range. *p*-values were calculated by Mann-Whitney test. (E) Percentage of animals expressing *grl-18*pro:YFP in zero, one, or two ILsoD glia in wild-type and *unc-130* mutant strains. Average number of cells marked per animal is listed under each condition. n = 50 animals per genotype. *p*-values were calculated by Fisher’s Exact test. (F) *unc-130* promoter was used to drive expression of *unc-130* and human *FOXD3* separately in the *unc-130* mutant strain and extent of rescue was assessed. Percentage of animals expressing *grl-18*pro:YFP in zero, one, or two ILsoD glia in each condition. Average number of cells marked per animal is listed under each condition. n = 50 animals per condition. *p*-values were calculated by Fisher’s Exact test.

From our mutant screen and previous studies, we assembled a collection of point mutations in the UNC-130 DBD: *ns313 -* W201*; oy10* and *ev659 -* R218*; op459 -* R219 (Nash et al., 2000; Sarafi-Reinach & Sengupta, 2000) (Fig. 5A, S6). Mapped onto the FoxD3 structure, W201 is located in a β-sheet that flanks the central helical bundle that inserts into the major groove of DNA, whereas the R218 and R219 residues are in a wing domain on the opposite side of the helix bundle (Fig. 5C) (Jin et al., 1999). To assess the effect of the W201G and R218C point mutations, we applied mutant UNC-130 recombinant proteins separately onto PBMs. We found that the R218C protein still preferentially binds DNA sequences recognized by wild-type UNC-130 but with lower affinity (Fig. 5D, S6), while the W201G mutant protein did not exhibit sequence-specific preferences for these or any other motifs, suggesting severely impaired DNA binding (Fig. 5D, S6). These binding defects strongly correlated with *in vivo* function. The W201G mutant exhibited defects in ILsoD specification that are nearly as strong as an early stop mutant (R62stop) or a deletion allele, such that >90% of ILsoD glia lose reporter expression (Fig. 5E). In contrast, in hypomorphic alleles, which include R218C, R218H, R219K, only 50% of ILsoD glia lose reporter expression (Fig. 5E). Thus, moderately impaired DNA binding appears to promote weak ILsoD glial defects, and severely impaired DNA binding results in strong ILsoD glial defects. We find that these highly conserved amino acid residues are crucial for UNC-130 function, and are likely to similarly disrupt FoxD3 function in vertebrates.

In vertebrates, FoxD3 is required for the specification of several cell types that arise from the neural crest, a multipotent lineage that participates in convergent differentiation (Dottori et al., 2001; Kos et al., 2001; Lister et al., 2006; Lukoseviciute et al., 2018; Sasai et al., 2001; Stewart et al., 2006; Teng et al., 2008). Although examination of UNC-130 and human FOXD3 protein sequences revealed little similarity outside of the highly-conserved DBD and eh1 motif (Fig. 5A, S6), we found that expression of an *unc-130* promoter::*FOXD3* cDNA transgene almost completely rescues ILsoD glial specification defects in *unc-130* mutants (Fig. 5F). This suggests that UNC-130 and FoxD3 functionality is conserved despite divergence at the primary sequence level.

In summary, although *C. elegans* lacks a neural crest, the similarities between FoxD3 and UNC-130 extend from their roles in lineage specification to their molecular mechanisms of action – including their preferred DNA binding sites and their roles as transcriptional repressors, likely via an interaction with the Groucho repressive complex. More broadly, our results lead to the speculation that prior to the evolution of the neural crest, there already existed a FoxD3 precursor that acted in a multipotent sublineage that exhibits convergent differentiation.

## DISCUSSION

Unraveling the mechanisms that establish fate competence in progenitors and precursors is critical to understanding how developmental lineage is coupled to cell fate – why do certain cell types arise only from particular lineages? The special case of convergent differentiation promises to offer important insights into this relationship.

An implication of our findings and previous studies is that different progenitors can take distinct transcriptional paths to produce the same cell type. We find that UNC-130 is only required in one of three convergent lineages to produce the ILso glial cell type. Although convergent differentiation was not their focus, previous studies provide additional examples. For example, loss of *mls-2/*Nkx affects mainly ventral, but not dorsal, CEPsh glia (Yoshimura et al., 2008). Similar to ILso glia, dorsal and ventral CEPsh glia are derived from lineages that diverge early in development. MLS-2 also appears to function at the lineage level, as other studies have determined that cell types related to CEPshV glia by lineage are mis-specified in *mls-2* mutants (Abdus-Saboor et al., 2012; Kim et al., 2010). Other examples include the requirement for LIN-32 to specify dorsal but not ventral CEP neurons, and CEH-10 to specify dorsal but not lateral or ventral RME neurons (Doitsidou et al., 2010; Forrester et al., 1998; Rojo Romanos et al., 2017). These observations provide tantalizing starting points for characterizing other factors that are involved in convergent differentiation.

Cell-type specification in terminally differentiating cells, which is mediated by master regulator transcription factors called terminal selectors, has been extensively studied in a number of cell types (Hobert & Kratsios, 2019). In contrast, factors that act transiently in progenitor cells to establish lineage-specific identity are less well-understood. Some candidates for this type of factor include CEH-36 and UNC-30, which function redundantly to regulate progenitor cell cycle progression and cell position in a number of developmental lineages (Walton et al., 2015), and CND-1, which regulates mitotic progression and establishment of neuronal fate in lineages that will give rise to neurons (Hallam et al., 2000). Another example is the lineage-restricted transcription factor TBX-37/38, which is expressed transiently early in embryogenesis to prime a locus that does not become active until several cell cycles later in the mature cell type (Charest et al., 2020; Cochella & Hobert, 2012). Similarly, in this study, we find that UNC-130, consistent with previous expression data from Sarafi-Reinach and Sengupta (2000), acts in lineage-related progenitor cells before these cells make terminal divisions and their progeny start to differentiate. Therefore, the function of these early-acting factors is likely to establish competency for fate specification, a role which differs from that of terminal selectors.

How does UNC-130 establish fate competency in a specific lineage? UNC-130 is required broadly for the specification of several unrelated cell types produced by the ILsoD sublineage. It is possible that UNC-130 primes and represses cell type-specific loci in each progenitor cell based on cellular context and binding partners. Alternatively, it may not impart any specific cell type information, but rather functions to transition progenitor cells to a more restricted state through the repression of pluripotency genes. Alternatively, or in addition, it may block off paths to alternate fates by repressing target genes that need to remain inactive in a particular lineage.

The results presented in this study are highly relevant to understanding the relationship between lineage and cell fate in the vertebrate nervous system, an intricate structure comprising many diverse cell types arising from a myriad of lineages. Single-cell RNA profiling of the mammalian brain has shown that the glial classes of astrocytes and microglia, which had long been thought to consist of molecularly homogeneous cells, actually exhibit striking region-specific molecular heterogeneity (Hammond et al., 2019; John Lin et al., 2017; Marques et al., 2016; Masuda et al., 2019; Morel et al., 2017; Spitzer et al., 2019; Zeisel et al., 2015, 2018). In contrast, the molecular signatures of oligodendrocytes are highly similar (Marques et al., 2018; Zeisel et al., 2018). Because astrocytes and oligodendrocytes are thought to share a common progenitor, these observations suggest that regionally distinct progenitor cells may undergo convergent differentiation in the mammalian nervous system as well. Elucidation of these pathways will require single-cell profiling methods to be combined with careful lineage tracing and functional perturbation of regulatory factors *in vivo* to ultimately achieve the level of resolution that is available in the far simpler nervous system of *C. elegans*.

## ACKNOWLEDGMENTS

We thank Laura Moriarty for assistance with the genetic screen that isolated *ns313*; Oliver Hobert for whole genome sequencing; Piali Sengupta for *unc-130* cDNA and the *oy10* mutant strain; Joseph Culotti for *unc-130* mutant strains; Raphael Bruckner for assistance with cell culture; Constance Cepko, Christopher Walsh, and the members of the Heiman laboratory for comments and advice on the manuscript; and WormBase. Some strains were provided by the CGC, which is funded by NIH Office of Research Infrastructure Programs (P40 OD010440).

## AUTHOR CONTRIBUTIONS

Conceptualization: K.M. and M.G.H.; Experimental design: K.M. and M.G.H.; PBM experiment and analysis: J.M.R.; Lineaging analysis: J.D.R. and J.I.M.; Writing of manuscript: K.M. and M.G.H. with input from all authors; Supervision and funding: S.S., M.L.B., J.I.M. and M.G.H.

## MATERIALS AND METHODS

### Strains

Strains were constructed in the N2 background and cultured under standard conditions (Brenner, 1974). Transgenic strains were generated with standard techniques (Mello & Fire, 1995) with injection of 100 ng/µL of DNA (5-50µL per plasmid). Strains, transgenes, and plasmids are listed in Tables S1-S3 respectively.

### Isolation and mapping of *unc-130* alleles

We isolated an allele of *unc-130, ns313,* from a genetic screen for sense organ abnormalities. Animals of genotype *oyIs44* V were mutagenized using 70 mM ethyl methanesulfonate (EMS, Sigma) at 20°C for 4 hours. Nonclonal F2 progeny were examined on a fluorescence stereomicroscope and animals with sense organ defects were recovered. A mutant strain, *ns313*, exhibiting short amphid dendrites (14% penetrance) and amphid sheath migration defects (55% penetrance) was isolated.

With standard linkage mapping and SNP analysis (Wicks et al., 2001), *ns313* was mapped to an interval between -6 cM and 15 cM on LG II. *ns313* animals were crossed to the Hawaiian strain CB4856 and F2 progeny with the mutant phenotype were transferred to individual plates. All F3 recombinants were pooled and subjected to genomic DNA extraction and whole-genome sequencing for one-step mapping (Doitsidou et al., 2010). Analysis with CloudMap (Minevich et al., 2012) identified a linked region on LG II including a point mutation in *unc-130* (GAACTAT[T>G]GGGCGTGGA) (W201G).

### Characterization of glial phenotypes

To score regional defects in IL sockets, we generated strains that co-expressed the ILso marker (*hmnIs47* [*grl-18*pro:mApple]) with a marker for URX (*ynIs48* [*flp-8*pro:GFP]), a dorsally located neuron whose dendrites fasciculate with the processes of the dorsal, but not lateral or ventral, ILso glia as a landmark. Glial specification defects were scored visually on either a Nikon SMZ1500 stereomicroscope with an HR Plan Apo 1.6x objective or a Deltavision Core imaging system (Applied Precision) with UApo/340 40x 1.35NA objective (Olympus).

### Fluorescence microscopy and image processing

Animals were mounted on 2% agarose pads in M9 buffer (Sulston et al., 1983) with 50-100 mM sodium azide depending on developmental stage, and imaged using a Deltavision Core imaging system (Applied Precision) with UApo/340 40x 1.35NA, PlanApo 60x 1.42NA, and U-PlanApo 100x 1.4NA objectives (Olympus) and CoolSnap HQ2 camera. Images were deconvolved using Softworx (Applied Precision) and maximum-brightness projections were obtained from contiguous optical sections using ImageJ.

### Lineaging analysis

Cell lineage analysis was performed using the StarryNite/AceTree cell tracking system (Bao et al., 2006; Boyle et al., 2006; Santella et al., 2010). Embryos from strains RW11144 [*ev505 ujIs113; wgIs476*] and CHB3933 [*ujIs113; wgIs746*] were imaged on a Leica SP5 resonance-scanning confocal microscope with approximately 1.5 min time point spacing and 0.5 micron z-resolution as previously described (Richards et al., 2013). Computational cell tracking of histone-mCherry images was used to track cells across time and identify the time of all terminal divisions in the lineages leading to ILsoD (ABp(l/r)aapa), ILsoL/R (ABalaaap) and ILsoV (ABalppap and ABarappp).

### Heat-shock induced expression

Strain CHB4158 [*ev505*; hmnEx2283 [*hsp16-2*pro:unc-130 + *hsp16-41*pro:unc-130 + *grl-18*pro:YFP + *unc-122*pro:RFP]] was grown at 20°C prior to heat-shock experiments. Age of embryos was determined by morphological state. Embryos at different developmental stages were sorted onto separate plates and heat-shocked at 34°C for 60 min to induce expression of UNC-130. After heat-shock, animals were grown at 20°C for 12-18 hours and embryos and larvae were scored for *grl-18*pro:YFP expression by fluorescence microscopy as described above.

### PBM experiments and data analysis

PBM experiments were performed on universal “all-10-mer” arrays in 8X60K format (Agilent, AMADID 030236) (Berger et al., 2006; Nakagawa et al., 2013). PBM experiments were performed at 500 nM protein concentration in the standard protein binding reaction mixture, substituting buffer A for PBS (buffer A= 138 mM KGlu, 12 mM NaHCO_3_, 0.8 mM MgCl_2_, pH 7.2) in the standard PBM protocol (Berger & Bulyk, 2009). Protein binding was detected with an Alexa488-conjugated anti-GST antibody (Life Technologies A-11131), and arrays were scanned using a GenePix 4400A (Molecular Devices) microarray scanner. Binding was quantified using the Universal PBM Analysis Suite (Berger & Bulyk, 2009) to generate E-scores for each 8-mer. Motifs were derived using the Seed-and-Wobble algorithm (Berger et al., 2006; Berger & Bulyk, 2009). Two replicate experiments were performed, with replicate 1 having higher E-scores overall. Replicate 1 is shown in Fig. 5D, and Replicate 2 is shown in Fig. S6B.

Boxplots were generated in R, from the E-scores of the 8-mer sequences that match the UNC-130 FkhP motif ([AG][CT]AAACA) or the FkhS motif (AA[CT]AACA). Individual data points are displayed on the boxplots using the stripchart function in R. Significant differences in binding were evaluated using a one-sided Mann-Whitney test, with the wilcox.test function in R. All PBM raw data are available on UniPROBE: http://thebrain.bwh.harvard.edu/uniprobe/ with deposition ID MIZ19A

### Luciferase assays

HEK293T cells were cultured at 37°C and transfected with Fugene (Roche). 48 h post transfection, cells were collected in cold 1X PBS and transferred into 96-well plates. Renilla and firefly luciferase activity were assayed according to manufacturer’s instructions using the Dual-Glo assay (Promega) and bioluminescence was collected on a Molecular Devices Spectramax Paradigm plate reader. Firefly luciferase activity was normalized to renilla luciferase activity in each sample.

