## Supplemental Material for "Lineage-specific control of convergent differentiation by a Forkhead repressor"

Figures S1-S6

Tables S1-S7

**Fig. S1**

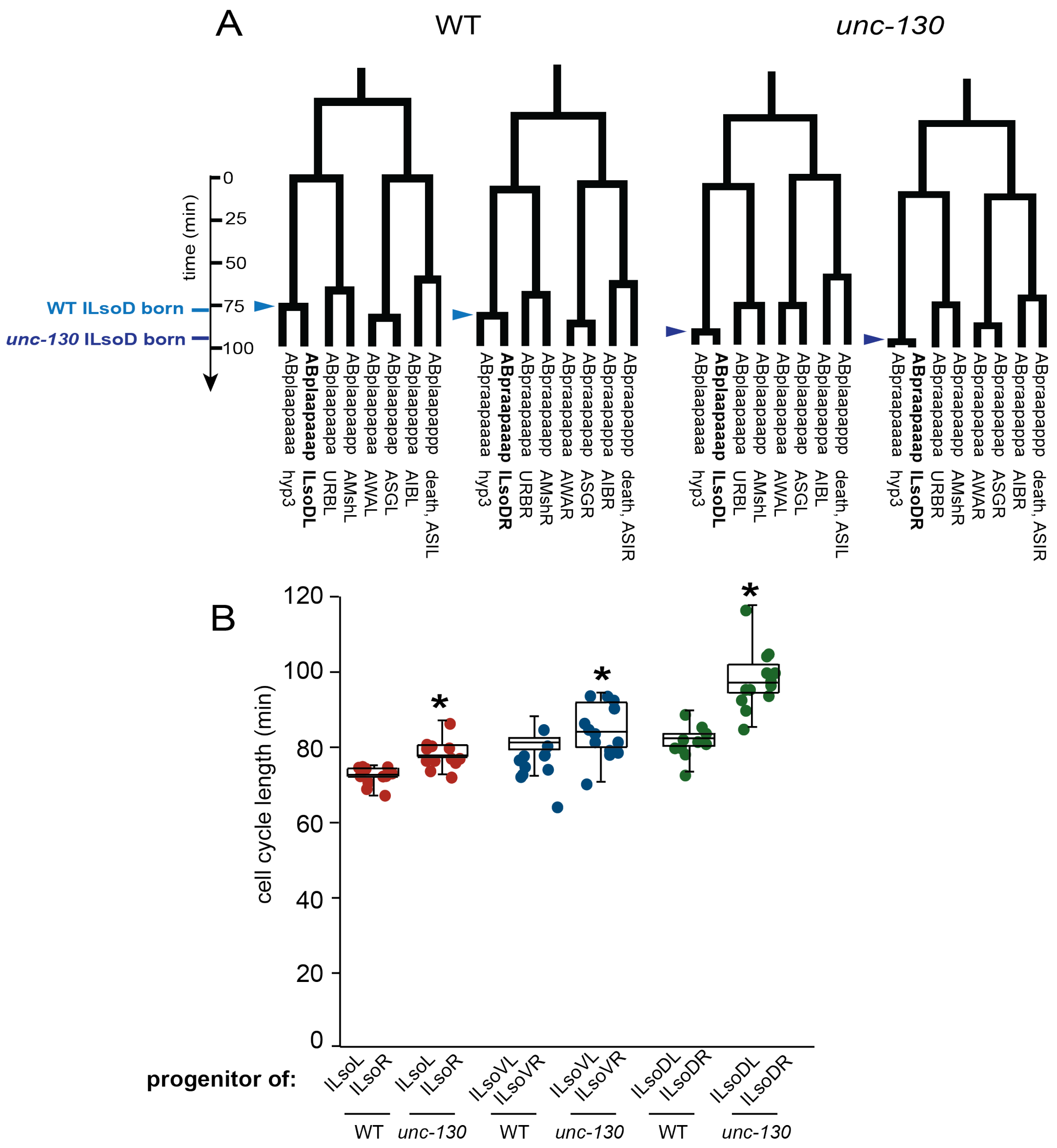
A
WT
unc-130
time (min)
0
25
50
75
100
WT ILsoD born
unc-130 ILsoD born
ABplaapaaaa hyp3
ABplaapaaap ILsoDL
ABplaapaapa URBL
ABplaapaapp AMshL
ABplaapapaa AWAL
ABplaapapap ASGL
ABplaapappa AIBL
ABplaapappp death, ASIL
ABpraapaaaa hyp3
ABpraapaaap ILsoDR
ABpraapaapa URBR
ABpraapaapp AMshR
ABpraapapaa AWAR
ABpraapapap ASGR
ABpraapappa AIBR
ABpraapappp death, ASIR
B
cell cycle length (min)
120
100
80
60
40
20
0
*
progenitor of:
ILsoL ILsoR
ILsoVL ILsoVR
ILsoDL ILsoDR
WT
unc-130

**Fig S1. Related to Fig. 1**

(A) Two example ILsoD (ABpxaapaa) lineages from wild-type and *unc-130* mutant embryos. Light blue arrowheads denote division of wild-type ILsoD progenitors and dark blue arrowheads denote divisions of *unc-130* mutant ILsoD progenitors that produce ILsoD and hyp3 cells. (B) Cell cycle lengths of ILsoL, ILsoR, ILsoVL, ILsoVR, ILsoDL, and ILsoDR progenitor cells in wild-type and *unc-130* mutant embryos. n = 6 for wild type, n = 8 for *unc-130* mutants. Center lines represent population median. Asterisks denote $p$-value < 0.05 as calculated by Welch's t-test.

**Fig. S2.**

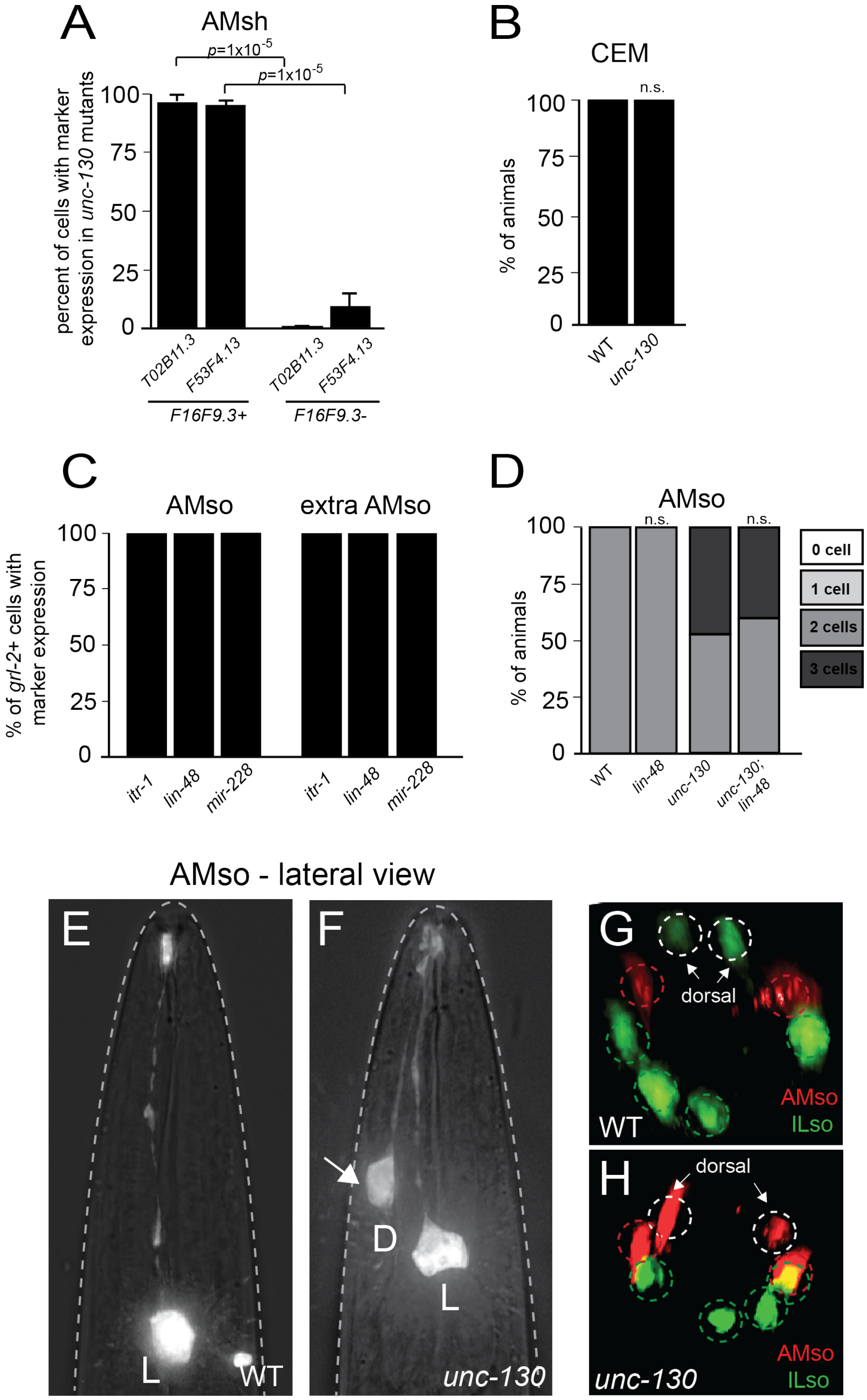

A
AMsh
p=1x10-5
p=1x10-5
percent of cells with marker expression in unc-130 mutants
100
75
50
25
0
T02B11.3
F53F4.13
T02B11.3
F53F4.13
F16F9.3+
F16F9.3-
B
CEM
n.s.
% of animals
100
75
50
25
0
WT
unc-130
C
AMso
extra AMso
% of grl-2+ cells with marker expression
100
75
50
25
0
itr-1
lin-48
mir-228
itr-1
lin-48
mir-228
D
AMso
n.s.
n.s.
% of animals
100
75
50
25
0
WT
lin-48
unc-130
unc-130; lin-48
0 cell
1 cell
2 cells
3 cells
AMso - lateral view
E
L
WT
F
D
L
unc-130
G
dorsal
WT
AMso
ILso
H
dorsal
unc-130
AMso
ILso

**Fig. S2. Related to Fig. 2**

(A) Percentage of *F16F9.3*+ and *F16F9.3-* cells co-expressing AMsh markers, *T02B11.3*pro:GFP and *F53F4.13*pro:GFP, in *unc-130* mutant animals. n ≥ 20 cells per marker. Error bars – standard error of proportion. *p*-values calculated by z-score for two population proportions. (B) Percentage of wild-type and *unc-130* mutant males expressing the CEM neuron marker, *pkd-2*pro:GFP. The CEM neuron is the sister cell of AMso in males. (C) Percentage of *grl-2*+ cells co-expressing AMso markers, *itr-1*pro:GFP and *lin-48*pro:GFP, and glial marker *mir-228*pro:histone-GFP in endogenous and extra cells in *unc-130* mutants. n ≥ 20 cells per marker. (D) Percentage of animals expressing AMso-specific marker *grl-2*pro:YFP in zero, one, two, three or more cells in wild type, *lin-48*, *unc-130*, and *lin-48*; *unc-130* double mutants. n=50 animals per genotype. *p*-values were calculated by Fisher's Exact test. Lateral views of wild-type (E) and *unc-130* mutant (F) animals expressing AMso marker, *grl-2*pro:YFP. Arrow indicates extra cell. In lateral views, the other endogenous AMso cell is not visible. D – dorsal, L – lateral. En face views of wild-type (G) and *unc-130* mutant (H) animals co-expressing *grl-18*pro:GFP to mark ILso glia and *grl-2*pro:mApple to mark AMso glia. Arrows indicate cells in the dorsal position. There are two extra AMso glia in this particular *unc-130* mutant animal.

**Fig S3.**

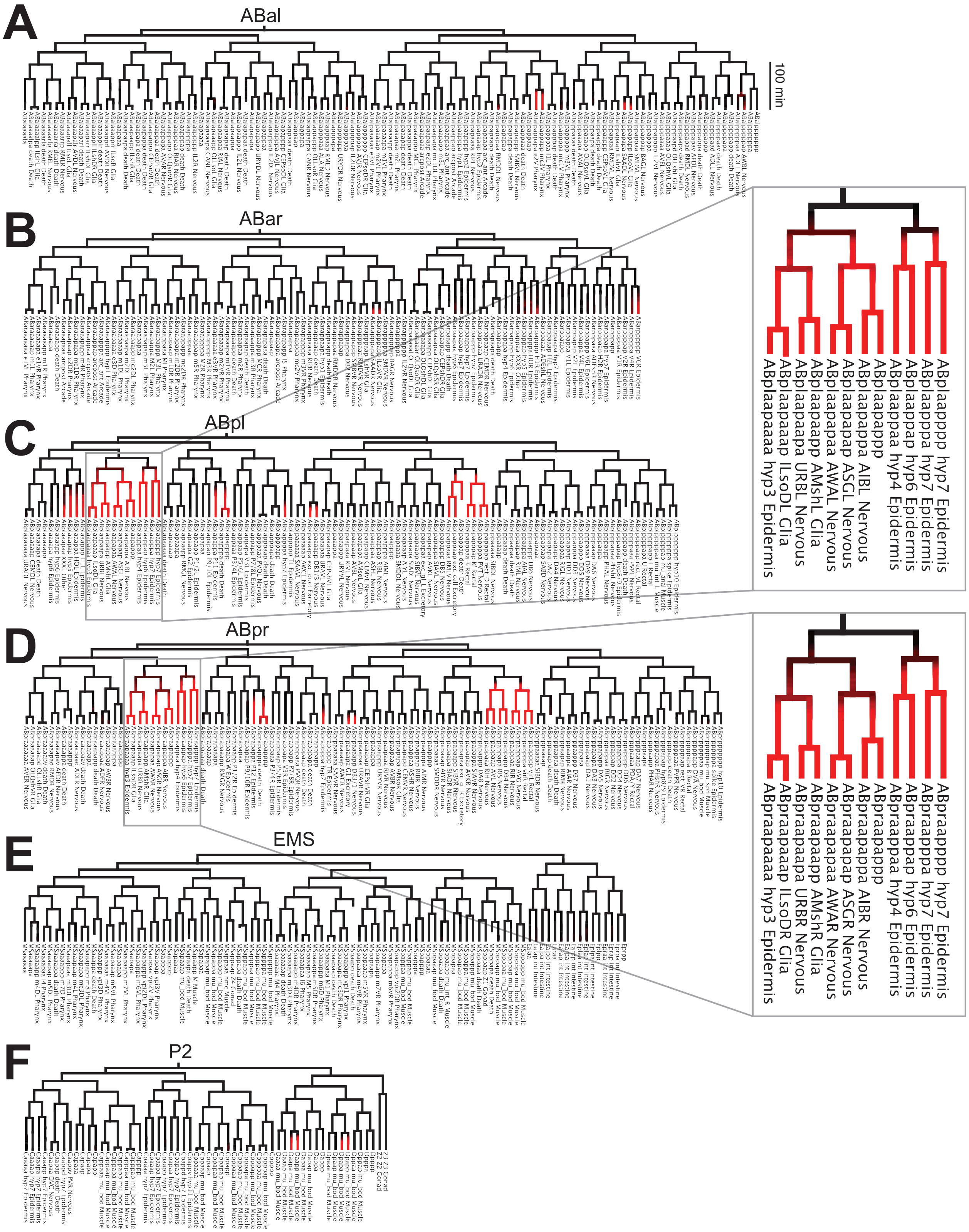
A
ABal
100 min
B
ABar
C
ABpl
D
ABpr
E
EMS
F
P2
ABplaapppp hyp7 Epidermis
ABplaapppa hyp7 Epidermis
ABplaappap hyp6 Epidermis
ABplaappaa hyp4 Epidermis
ABplaapapp
ABplaapappa AIBL Nervous
ABplaapapap ASGL Nervous
ABplaapapaa AWAL Nervous
ABplaapaapp AMshL Glia
ABplaapaapa URBL Nervous
ABplaapaaap ILsoDL Glia
ABplaapaaaa hyp3 Epidermis
ABpraapppp hyp7 Epidermis
ABpraapppa hyp7 Epidermis
ABpraappap hyp6 Epidermis
ABpraappaa hyp4 Epidermis
ABpraapapp
ABpraapappa AIBR Nervous
ABpraapapap ASGR Nervous
ABpraapapaa AWAR Nervous
ABpraapaapp AMshR Glia
ABpraapaapa URBR Nervous
ABpraapaaap ILsoDR Glia
ABpraapaaaa hyp3 Epidermis

**Fig. S3. Related to Fig. 3**

(A-F) Lineage diagram of all embryonic divisions in a wild-type embryo from the lineaging strain RW11144, which expresses UNC-130:GFP. (A) ABal, (B) ABar, (C) ABpl, (D) ABpr, (E) MS and E, and (F) C, D, and Z lineages.  UNC-130 expression levels are indicated in red.

**Fig. S4**

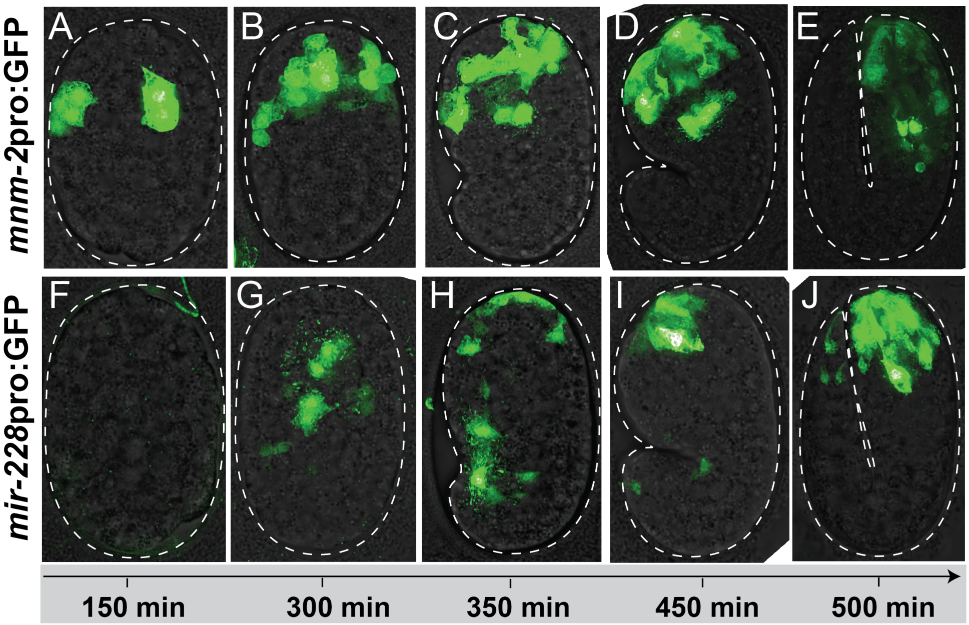


**Figure S4. Related to Fig. 3.**

Time course of *mnm-2*pro:GFP (A-E) and *mir-228*pro:GFP (F-J) expression in embryos.

**Fig S5.**

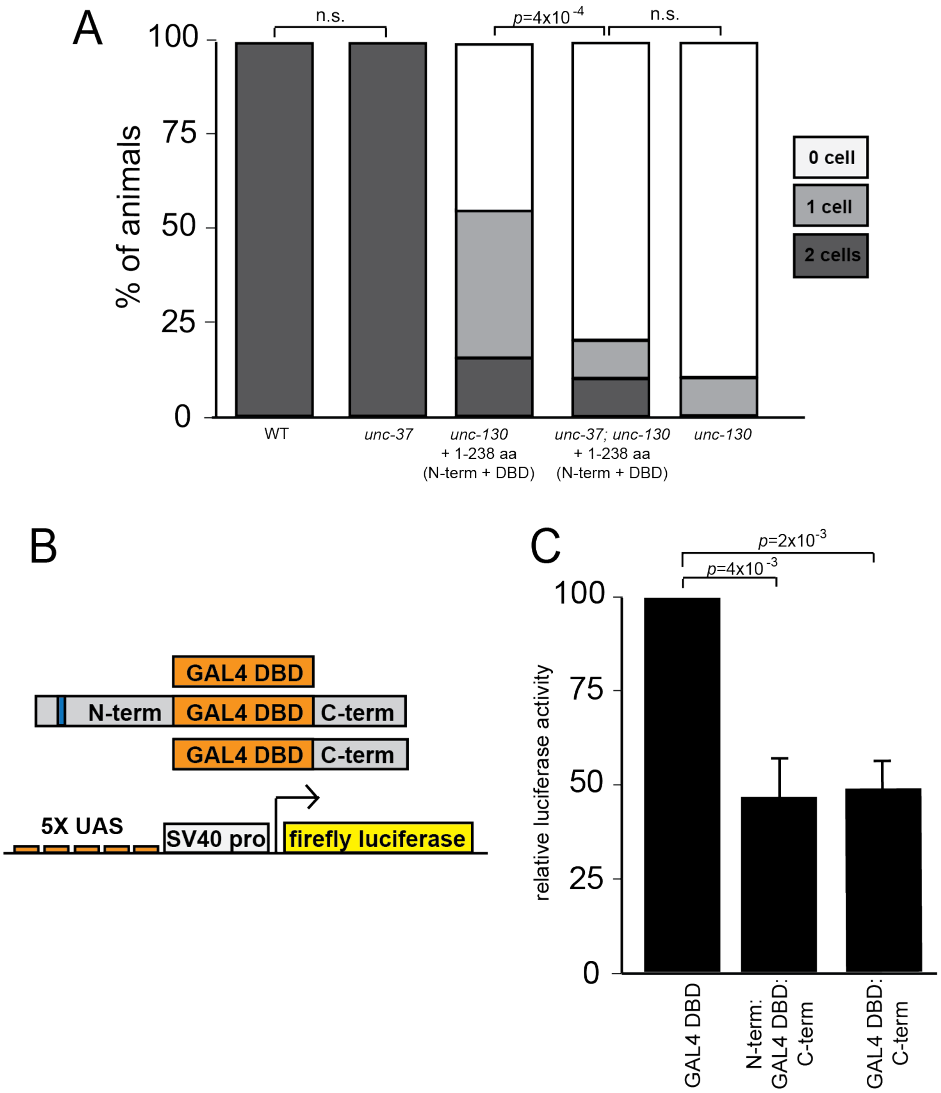


**Fig S5. Related to Fig. 4**

(A) Percentage of animals expressing *grl-18*pro:YFP in zero, one, or two ILsoD glia in wild-type, *unc-37* mutants, and *unc-130* mutants with N-terminus rescue, *unc-130*; *unc-37* double mutants with N-terminus rescue, and *unc-130* mutants alone. n = 50 animals per condition. *p*-values were calculated by Fisher's Exact test. (B) Schematic diagram of constructs used in luciferase assays. (C) GAL4DBD alone, N-term:GAL4DBD:C-term, or GAL4DBD:C-term, along with a UAS-firefly luciferase reporter and constitutively expressed renilla luciferase were transfected into HEK293T cells. Relative firefly luciferase activity, first normalized to renilla luciferase bioluminescence in each sample, and then to DBD-GAL4 alone. Error bars - SD; *p*-values calculated by Welch's t-test with Bonferroni correction for multiple t-tests.

**Fig. S6**

A

```
UNC-130 ---------MLFSMESILSSTKPKLEPPPKLEP--EVTINEQVVDLPR--SNTRLSEPST
FoxD3   MTLSGGGSASDMSGQTVLTAEDVDIDVVGEGDDGLEEKDSDAGCDSPAGPPELRLDEADE

        SASVLEHDLKFGESRKRSRSLGDEPTEDEDGVPVRKANKRNHSTSSAADSSSDDAKDDDD
        VPPAAPHHGQPQPPHQQPLTLPKEAAGAGAGPGGDVGAPEADGCKGGVGGEEGGASGGGP

        DDDSTSRKSMSGHR-KSSHAKPPYSYIALIAMSILNSPEKKLTLSEICEFIINKFEYYKE
        GAGSGSAGGLAPSKPKNSLVKPPYSYIALITMAILQSPQKKLTLSGICEFISNRFPYYRE
                            ********** * ** ** ****** ***** * **** *

        KFPAWQNSIRHNLSLNDCFVKVARGPGNPGKGNYWALDPNCEDMFDNGSFLRRRKRYKKN
        KFPAWQNSIRHNLSLNDCFVKIPREPGNPGKGNYWTLDPQSEDMFDNGSFLRRRKRFKRH
        *********************    ********** ***   *************** *

        SDTYH------EMMSHHPMPFPPFLPQGMPFP-PRMMHPMANIPMLGHPMNPRAVPNMPA
        QQEHLREQTALMMQSFGAYSLAAAAGAAGPYGRPYGLHPAAAAFAYSHPAAAAAAAAAAA

        FFIPQNID--------------------------------SQKLLSMMASRIMPMDAPVS
        LQYPYALPPVAPVLPPAVPLLPSGELGRKAAAFGSQLGPGLQLGLNSLGAAAAAAGTAGA

        SGQKRTSSSSSSPNENGSSAVSDKLSA---------------------------------
        AGTTASLIKSEPSARPSFSINIIGGGPAAPGGSAVGAGVAGGTGGSGGGSTAQSFLRPP
        ------------------------------------------------------------
        GTVQSAALMATHQPLSLSRTTATIAPILSVPLSGQFLQPAASAAAAAAAAAQAKWPAQ
```

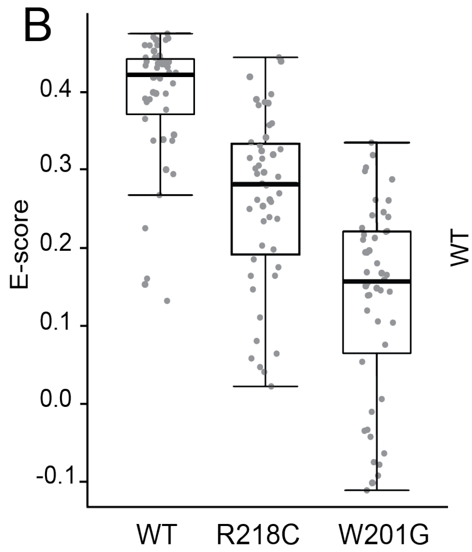


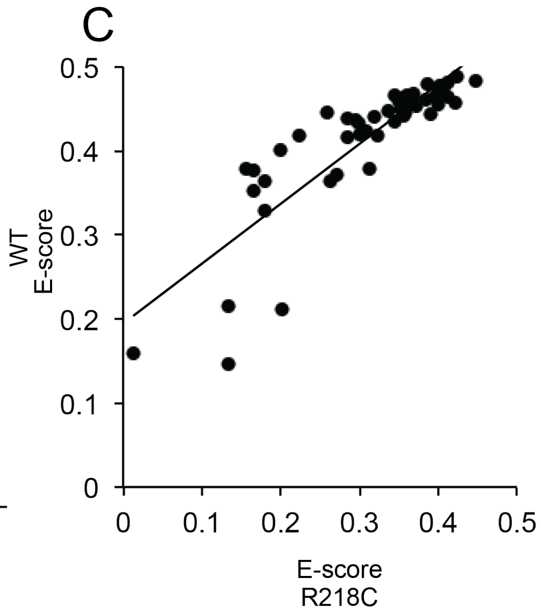

**Fig. S6. Related to Fig. 5**

(A) Alignment of UNC-130 (black) and FoxD3 (gray) sequences. DNA binding domain is highlighted in green. Eh1 motifs are highlighted in blue. Asterisks denote conserved amino acids in the DNA binding domain. Underlined residues are mutated in point mutants described in Fig. 5. (B) Technical replicate of PBM experiment from Fig. 5D. Scatter plot of E-scores for 8-mer DNA sequences matching [A/G][C/T]AAACA or AA[C/T]AACA from protein binding microarray (PBM) assays of wild-type, R218C, and W201G mutant proteins. Black lines represent population median, top and bottom of boxes are 25$^{th}$ and 75$^{th}$ percentiles, respectively, and top and bottom of whiskers are either most extreme point or 1.5x the interquartile range. *p*-values calculated by Mann-Whitney test. (C) Scatter plot of E-scores for 8-mer DNA sequences matching [A/G][C/T]AAACA or AA[C/T]AACA for wild-type versus R218C mutant proteins. Linear regression, $R^2$=0.72

**Table S1. Penetrance of phenotypes in Fig. 1**

| | | | % of animals with # of cells | | | | |
|---|---|---|---|---|---|---|---|
| **cell type** | **marker** | **genotype** | **0** | **1** | **2** | **n** | ***p*-value** |
| **ILsoD** | *grl-18* | wt | 0 | 0 | 100 | 50 | $2x10^{-29}$ |
| | | *ev505* | 90 | 10 | 0 | 50 | |
| **ILsoD** | *col-53* | wt | 0 | 0 | 100 | 50 | $2x10^{-29}$ |
| | | *ev505* | 88 | 12 | 0 | 50 | |
| **ILsoD** | *col-177* | wt | 0 | 6 | 94 | 50 | $2x10^{-27}$ |
| | | *ev505* | 86 | 14 | 0 | 50 | |
| **ILsoL/R** | *grl-18* | wt | 0 | 0 | 100 | 50 | 1 |
| | | *ev505* | 0 | 0 | 100 | 50 | |
| **ILsoL/R** | *col-53* | wt | 0 | 6 | 94 | 50 | 0.1 |
| | | *ev505* | 0 | 0 | 100 | 50 | |
| **ILsoL/R** | *col-177* | wt | 8 | 10 | 82 | 50 | 0.4 |
| | | *ev505* | 2 | 14 | 84 | 50 | |
| **ILsoV** | *grl-18* | wt | 0 | 0 | 100 | 50 | 1 |
| | | *ev505* | 0 | 0 | 100 | 50 | |
| **ILsoV** | *col-53* | wt | 0 | 4 | 96 | 50 | 1 |
| | | *ev505* | 0 | 4 | 96 | 50 | |
| **ILsoV** | *col-177* | wt | 0 | 2 | 98 | 50 | 0.1 |
| | | *ev505* | 0 | 12 | 88 | 50 | |

**Table S2. Penetrance of IL neuron phenotypes.**

| | | | % of animals with # of cells | | | | | |
|---|---|---|---|---|---|---|---|---|
| **cell type** | **marker** | **genotype** | **4** | **5** | **6** | **>6** | **n** | ***p*-value** |
| **IL1** | *flp-3* | wt | 12 | 10 | 78 | 0 | 50 | $9x10^{-4}$ |
| | | *ev505* | 16 | 4 | 58 | 22 | 50 | |
| **IL2** | *klp-6* | wt | 4 | 12 | 84 | 0 | 50 | 0.4 |
| | | *ev505* | 0 | 12 | 84 | 4 | 50 | |

**Table S3. Penetrance of phenotypes in Fig. 2.**

| | | | % of animals with # of cells | | | | | | |
|---|---|---|---|---|---|---|---|---|---|
| **cell type** | **marker** | **genotype** | **0** | **1** | **2** | **3** | **4** | **n** | ***p*-value** |
| **hyp3** | *ceh-10* | wt | 0 | 0 | 100 | - | - | 50 | $1x10^{-10}$ |
| | | *ev505* | 0 | 30 | 70 | - | - | 50 | |
| **URB** | *nlp-6* | wt | 0 | 2 | 98 | - | - | 50 | 1 |
| | | *ev505* | 0 | 4 | 96 | - | - | 50 | |
| **AMsh** | *F16F9.3* | wt | 0 | 0 | 100 | - | - | 50 | $5x10^{-7}$ |
| | | *ev505* | 6 | 32 | 62 | - | - | 53 | |
| **CEPsh** | *hlh-17* | wt | 0 | 0 | 0 | 0 | 100 | 50 | 0.2 |
| | | *ev505* | 0 | 0 | 0 | 6 | 94 | 50 | |
| **AMso** | *grl-2* | wt | 0 | 0 | 100 | 0 | 0 | 50 | $7x10^{-8}$ |
| | | *ev505* | 0 | 0 | 58 | 34 | 8 | 53 | |
| **PHsh** | *F16F9.3* | wt | 0 | 12 | 88 | - | - | 50 | 1 |
| | | *ev505* | 0 | 14 | 86 | - | - | 50 | |
| **PHso** | *grl-2* | wt | 0 | 0 | 0 | 4 | 96 | 50 | 0.2 |
| | | *ev505* | 0 | 0 | 0 | 12 | 88 | 50 | |

**Table S4. Strains generated for this study**

| ID | Genotype | Figures |
|---|---|---|
| CHB3747 | *hmnEx2123* [*grl-2*pro:CFP + *grl-18*pro:YFP] | 1 |
| CHB3756 | *unc-130 (ev505); hmnEx2126* [*grl-2*pro:CFP + *grl-18*pro:YFP] | 1 |
| CHB3310 | *hmnIs47* [*grl-18*pro:mApple]*; ynIs78* [*flp-8*pro:GFP] | 1, 5 |
| CHB3311 | *unc-130 (ev505); hmnIs47* [*grl-18*pro:mApple]*; ynIs78* [*flp-8*pro:GFP] | 1 |
| CHB4124 | *hmnIs47* [*grl-18*pro:mApple]; *hmnEx2227* [*col-53*pro:GFP + pRF4] | 1 |
| CHB4143 | *unc-130 (ev505); hmnIs47* [*grl-18*pro:mApple]; *hmnEx2227* [*col-53*pro:GFP + pRF4] | 1 |
| CHB4064 | *hmnIs47* [*grl-18*pro:mApple]; *hmnEx2171* [*col-177*pro:GFP + pRF4] | 1 |
| CHB4125 | *unc-130 (ev505); hmnIs47* [*grl-18*pro:mApple]; *hmnEx2171* [*col-177*pro:GFP + pRF4] | 1 |
| CHB3933 | *ujIs113; wgIs476* | S1 |
| CHB4067 | *ev505 ujIs113; wgIs476* | S1 |
| CHB4046 | *hmnIs100* [*ceh-10*pro:GFP + pRF4] | 2 |
| CHB4047 | *unc-130 (ev505); hmnIs100* [*ceh-10*pro:GFP + pRF4] | 2 |
| CHB4066 | *hmnEx2237* [*nlp-6*pro:GFP + pRF4] | 2 |
| CHB4163 | *unc-130 (ev505); hmnEx2237* [*nlp-6*pro:GFP + pRF4] | 2 |
| CHB3562 | *unc-130 (ev505)*; *irIs67* [*hlh-17*pro:GFP] | 2 |
| CHB1634 | *unc-130 (ev505); hmnIs13* [*F16F9.*3pro:mCherry + *grl-2*pro:YFP + *gcy-8*pro:CFP] | 2 |
| CHB1549 | *hmnIs13* [*F16F9.3*pro:mCherry + *grl-2*pro:YFP + *gcy-8*pro:CFP] | 2 |
| CHB3850 | *hmnEx1910* [*grl-2*pro:mCherry + *itr-1*pro:YFP + pRF4] | S2 |
| CHB3355 | *unc-130 (ev505); hmnEx1910* [*grl-2*pro:mCherry + *itr-1*pro:YFP + pRF4] | S2 |
| CHB3422 | *saIs14* [*lin-48*pro:GFP]; *hmnEx1939* [*grl-2*pro:mCherry + pRF4] | S2 |
| CHB3441 | *unc-130 (ev505); saIs14* [*lin-48*pro:GFP]; *hmnEx1951* [*grl-2*pro:mCherry; pRF4] | S2 |

| | | |
|---|---|---|
| CHB3221 | *hmnEx1715* [*F53F4.13*pro:GFP + pRF4]*; hmnIs13* [*F16F9.*3pro:mCherry + *grl-2*pro:YFP + *gcy-8*pro:CFP] | S2 |
| CHB2966 | *hmnEx1683* [*T02B11.3*pro:GFP + pRF4]*; hmnIs13* [*F16F9.*3pro:mCherry + *grl-2*pro:YFP + *gcy-8*pro:CFP] | S2 |
| CHB3045 | *unc-130 (ev505); hmnEx1715* [*F53F4.13*pro:GFP + pRF4]*; hmnIs13* [*F16F9.*3pro:mCherry + *grl-2*pro:YFP + *gcy-8*pro:CFP] | S2 |
| CHB3030 | *unc-130 (ev505); hmnEx1683* [*T02B11.3*pro:GFP + pRF4]*; hmnIs13* [*F16F9.*3pro:mCherry + *grl-2*pro:YFP + *gcy-8*pro:CFP] | S2 |
| CHB2223 | *unc-130 (ev505); myIs4* [*pkd-2*pro:GFP] *him-5* (*e1490*) | S2 |
| CHB4308 | *unc-130 (ev505); hmnEx2293* [*mir-228*pro:histone-GFP]; *hmnEx1949* [*grl-2*pro:mApple + pRF4] | S2 |
| CHB4309 | *unc-130 (ev505); lin-48 (sa469); hmnIs13* [*F16F9.*3pro:mCherry + *grl-2*pro:YFP + *gcy-8*pro:CFP] | S2 |
| CHB4302 | *lin-48 (sa469); hmnIs13* [*F16F9.*3pro:mCherry + *grl-2*pro:YFP + *gcy-8*pro:CFP] | S2 |
| CHB2958 | *hmnEx1676* [*grl-2*pro:mApple + *grl-18*pro:GFP] | S2 |
| CHB2961 | *unc-130 (ev505); hmnEx1679* [*grl-2*pro:mApple + *grl-18*pro:GFP] | S2 |
| CHB3775 | *hmnIs82* [*grl-18*pro:GFP] | 3 |
| CHB4160 | *unc-130 (ev505); hmnIs82* [*grl-18*pro*:*GFP] | 3 |
| CHB3313 | *unc-130 (ev505); hmnEx1903* [*unc-130*pro4:unc-130 + *grl-18*pro:GFP + *flp-8*pro:mCherry + pRF4] | 3, 4, 5 |
| CHB4174 | *unc-130 (ev505); hmnIs82* [*grl-18*pro*:*GFP]*; hmnEx2273* [*mnm-2*pro:unc-130 + pRF4] | 3 |
| CHB4157 | *unc-130 (ev505); hmnIs82; hmnEx2282* [*mir-228*pro:unc-130 + *grl-18*pro:YFP + pRF4] | 3 |
| CHB4158 | *unc-130 (ev505); hmnEx2283* [*hsp16-2*pro*:*unc-130 + *hsp16-41*pro*:*unc-130 + *grl-18*p:YFP + *unc-122*pro:RFP) | 3 |
| CHB1996 | *hmnEx1138* [*mnm-2*pro:GFP + *egl-38*pro:nls-mCherry + pRF4] | S4 |
| CHB3447 | *unc-130 (ev505); hmnEx1964* [*unc-130*pro4:DBD + *grl-18*pro:GFP + *flp-8*:mCherry + pRF4] | 4 |
| CHB3428 | *unc-130 (ev505); hmnEx1945* [*unc-130*pro4:DBD-VP64 + *grl-18*pro:GFP + *flp-8*:mCherry + pRF4] | 4 |
| CHB3381 | *unc-130 (ev505); hmnEx1922* [*unc-130*pro4:N-TERM + *grl-18*pro:GFP + *flp-8*:mCherry + pRF4] | 4 |

| | | |
|---|---|---|
| CHB3402 | *unc-130 (ev505); hmnEx1927* [*unc-130*pro4: del eh1 N-TERM + *grl-18*pro:GFP + *flp-8*:mCherry + pRF4] | 4 |
| CHB3427 | *unc-130 (ev505); hmnEx1944* [*unc-130*pro4:C-TERM + *grl-18*pro:GFP + *flp-8*:mCherry + pRF4] | 4 |
| CHB4144 | *unc-130 (ev505); hmnEx2275* [*unc-130*pro4:DBD-Engrailed + *grl-18*pro:GFP + pRF4] | 4 |
| CHB4139 | *unc-130 (ev505); hmnEx2274* [*unc-130*pro4: del eh1 unc-130 + *grl-18*pro:YFP + *F16F9.3*pro:CFP + pRF4) | 4 |
| CHB4311 | *unc-130 (ev505); unc-37(e262) hmnEx2345* [*unc-130*pro4:N-TERM + *grl-18*pro:GFP + *unc-122*pro:RFP] | S5 |
| CHB4310 | *unc-37 (e262); hmnIs82* [*grl-18*pro:GFP] | S5 |
| CHB3381 | *unc-130 (hd12); hmnIs47* [*grl-18*pro:mApple]*; ynIs48* [*flp-8*pro:GFP] | 5 |
| CHB3382 | *unc-130 (ns313); hmnIs47* [*grl-18*pro:mApple]*; ynIs48* [*flp-8*pro:GFP] | 5 |
| CHB3384 | *unc-130 (oy10); hmnIs47* [*grl-18*pro:mApple]*; ynIs48* [*flp-8*pro:GFP] | 5 |
| CHB3382 | *unc-130 (ev659); hmnIs47* [*grl-18*pro:mApple]*; ynIs48* [*flp-8*pro:GFP] | 5 |
| CHB3378 | *unc-130 (op459); hmnIs47* [*grl-18*pro:mApple]*; ynIs48* [*flp-8*pro:GFP] | 5 |
| CHB3411 | *unc-130 (ev505); hmnEx1936* [*unc-130*pro4:FOXD3, *grl-18*pro:GFP, *flp-8*:mCherry, pRF4] | 5 |

**Table S5. Strains generated in previous studies**

| **ID** | **Genotype** | **Fig.** | **Source/Reference** |
|---|---|---|---|
| OP476 | *wgIs476* [unc-86::TY1::EGFP::3xFLAG] | S1 | (Sarov et al., 2006) |
| VPR839 | *irIs67* [*hlh-17*pro:GFP]; *unc-119* | 2 | (Stout & Parpura, 2011) |
| CM117 | *saIs14* [*lin-48*pro:GFP] | S2 | (Johnson et al., 2001) |
| OP77 | *wgIs77* [unc-130::TY1::EGFP::3xFLAG] | 3 | (Sarov et al., 2006) |
| RW11144 | *itIs37* [*pie-1*pro:mCherry], *stIs10116* [*his-72*pro:his-24:mCherry]; *wgIs76* [unc-130:TY1:EGFP:3xFLAG] | 3, S3 | (Murray et al., 2012) |

| | | | |
|---|---|---|---|
| OS4260 | *nsIs198* [*mir-228*pro:GFP] | S4 | (Pierce et al., 2008) |
| PT621 | *myIs4* [*pkd-2*pro:GFP] *him-5* (*e1490*) | S2 | (Barr & Sternberg, 1999) |

**Table S6. Plasmids generated for this study**

| **ID** | **Name** |
|---|---|
| pKM47 | *grl-18*pro:YFP |
| pKM15 | *grl-2*pro:YFP |
| pKM117 | *grl-2*pro:CFP |
| pIL36 | *grl-2*pro:mCherry |
| pIL41 | *grl-2*pro:mApple |
| pKM16 | *F16F9.3*pro:CFP |
| pKM55 | pDest15 UNC-130 DBD |
| pKM69 | pDest15 R218C UNC-130 DBD |
| pKM71 | pDest15 W201G UNC-130 DBD |
| pKM59 | *unc-130*pro4:UNC-130 |
| pKM77 | *unc-130*pro4:N-term:UNC-130 DBD |
| pKM79 | *unc-130*pro4:UNC-130 DBD:C-term |
| pKM88 | *unc-130*pro4:del eh1 N-term:UNC-130 DBD |
| pKM78 | *unc-130*pro4:UNC-130 DBD |
| pKM83 | *unc-130*pro4:UNC-130 DBD VP64 |
| pKM80 | *unc-130*pro4: del eh1 UNC-130 |
| pKM115 | 5XUAS SV40pro:firefly luciferase |
| pKM114 | CAGpro: renilla luciferase |

| | |
|---|---|
| pKM111 | CAGpro:GAL4DBD:UNC-130 C-term |
| pKM113 | CAGpro:GAL4DBD |
| pKM108 | CAGpro:UNC-130 N-term:GAL4DBD:<br>UNC-130 C-term |
| pKM72 | *unc-130*pro4:FOXD3 |
| pKM119 | *mir-228*pro:unc-130 |
| pKM118 | *mnm-2*pro:unc-130 |
| pKM67 | *hsp16-2*pro:unc-130 |
| pKM68 | *hsp16-41*pro:unc-130 |
| pKM126 | *nlp-6*pro:GFP |
| pKM123 | *ceh-10*pro:GFP |

**Table S7. Primers of general interest**

| **Name** | **Sequence** |
|---|---|
| unc-130pro4_fwd | gtactCCTGCAGGctttcaattgaaaattccgaga |
| unc-130pro4_rev | gatcGGCGCGCCtgttACCGGTgtctacctagt |
